## Supplementary files for "AgRP1 modulates breeding season-dependent feeding behavior in female medaka"

```
#Copyright Kimura Tomoki & Shinji Kanda @ Dept.Biol.Sci, Grad.Sch.Sci, The University of Tokyo
#This script can be run under python 3.5. OpenCV3 should be installed.
#a key add the total number
#z key reset the total number
#q key quit the program
#space key pauses the program
#this script can be run with 3 parameters like python thisfilename.py 5 3 1

import sys
import numpy as np
import cv2
import time
def conv(frame, argvs):
	gray = cv2.cvtColor(frame, cv2.COLOR_BGR2GRAY)
	#binarization.INV means black and white inversion. blockSize means pixel cutoff.  3, 5, 7,etc.
	th1 = cv2.adaptiveThreshold(gray,255,cv2.ADAPTIVE_THRESH_GAUSSIAN_C,¥
			cv2.THRESH_BINARY_INV,29,8)
	#iterations – number of times dilation is applied.anchor – anchor of the kernel that indicates the relative position of a filtered point within the kernel; the anchor should lie within the kernel; default value (-1,-1) means that the anchor is at the kernel center.
	kernel = np.ones((int(argvs[1]),int(argvs[1])),np.uint8)
	dilation = cv2.dilate(th1, kernel, iterations =1)
	erosion = cv2.erode(dilation,kernel,iterations = int(argvs[2]))
	dilation = cv2.dilate(erosion, kernel, iterations =int(argvs[3]))
	return dilation
#draw a circle after detemining the center of each shape
def CoG(contours,frame):
	arr=[]
	for i in contours:
		x_=0
		y_=0
		for j in i:
			x_ += j[0][0]
			y_ += j[0][1]
		arr.append([int(x_/len(i)), int(y_/len(i))])
	arr = np.array(arr)
	for i in arr:
		 cv2.circle(frame,tuple(i), 5, (0,255,0), 1)
def count(frame,gray):
	#edge detection
	img, contours, hierarchy = cv2.findContours(gray,cv2.RETR_EXTERNAL,cv2.CHAIN_APPROX_SIMPLE)
```

```
	CoG(contours,frame)
	#img = cv2.drawContours(frame, contours, -1, (0,255,0), 1)

	return contours
totalNum = 0
lastLog = 0
pressCount = 0
argvs = sys.argv
argc = len(argvs)
if (argc != 4):
	print ('Usage: python count.py [kernel] [erosion] [dilation]¥nkernel...Increase the parameter to reduce the noize. It increases the possiblity to misditection of the object')
	print('erosion...Increase this value when you want to distinguish the objects conted as one.¥ndilation...Increase this value when one object is double counted. Too large value may cause multiple objects counted as one')
	quit()		# Program End
cap = cv2.VideoCapture(0) #0 is the number of the camera

print('fps={0:2d}, w={1}, h={2}'.format(int(cap.get(cv2.CAP_PROP_FPS)),cap.get(3), cap.get(4)))
print('kernel={0}, iterations of erosion={1}, iterations of dilatioin={2}'.format(int(argvs[1]),int(argvs[2]),int(argvs[3])))
while(cap.isOpened()):
	ret, frame = cap.read()

	if ret:
		fin_img=conv(frame, argvs)
		contours=count(frame,fin_img)	#Edge detection,  draw circles
		cv2.putText(frame,"realtime#="+str(len(contours)),(10,40), cv2.FONT_HERSHEY_COMPLEX, 1.0,(0, 0, 255))	#print number of object
		descrip = "space:pause q:quit"
		cv2.putText(frame,"space:pause q:quit a:add z:reset",(10,440), cv2.FONT_HERSHEY_COMPLEX, 0.5,(0, 255, 0))	#print number of object
		cv2.putText(frame,"total#=" + str(totalNum) + "(" + str(pressCount) + ")",(10,80), cv2.FONT_HERSHEY_COMPLEX, 1.0,(0, 0, 255))	#total number
		cv2.imshow('frame',frame)	#Out put 1 frame in the window
	else :
		break
	keycode = cv2.waitKey(1)	#wait 1ms for key input
	if keycode == ord('q'): #when q is pressed, program ends
		break
	elif keycode == ord(' '):	#when space key was pressed
		while(True):
			keycode = cv2.waitKey(0)	#wait key press forever
```

```
            if(keycode== ord(' ')):     #space key: pause
                break
            elif(keycode== ord('q')): #q key stops the program
                cap.release()
                out.release()
                cv2.destroyAllWindows()
                quit()
    elif keycode== ord('a'): #when a key is pressed,add the number to the total number
        if time.time() > lastLog+1: # avoid multiple count in a short time
            totalNum = len(contours) + totalNum
            lastLog = time.time()
            pressCount = pressCount+1

    elif keycode== ord('z'): #when z key is pressed, reset the total number and press count.
            totalNum = 0
            pressCount = 0


cap.release()
cv2.destroyAllWindows()
```

**Figure 1-Source code 1: The code of “Shrimp-counter” system**

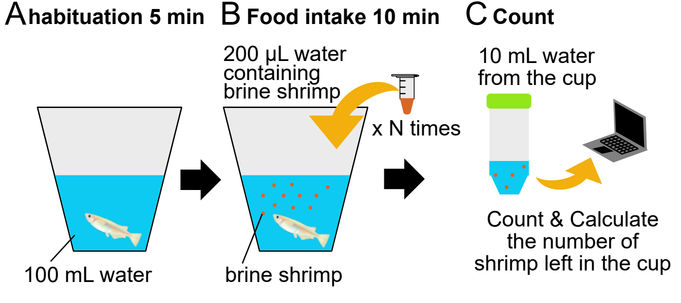
A habituation 5 min
B Food intake 10 min
C Count
200 µL water
containing
brine shrimp
x N times
10 mL water
from the cup
100 mL water
brine shrimp
Count & Calculate
the number of
shrimp left in the cup

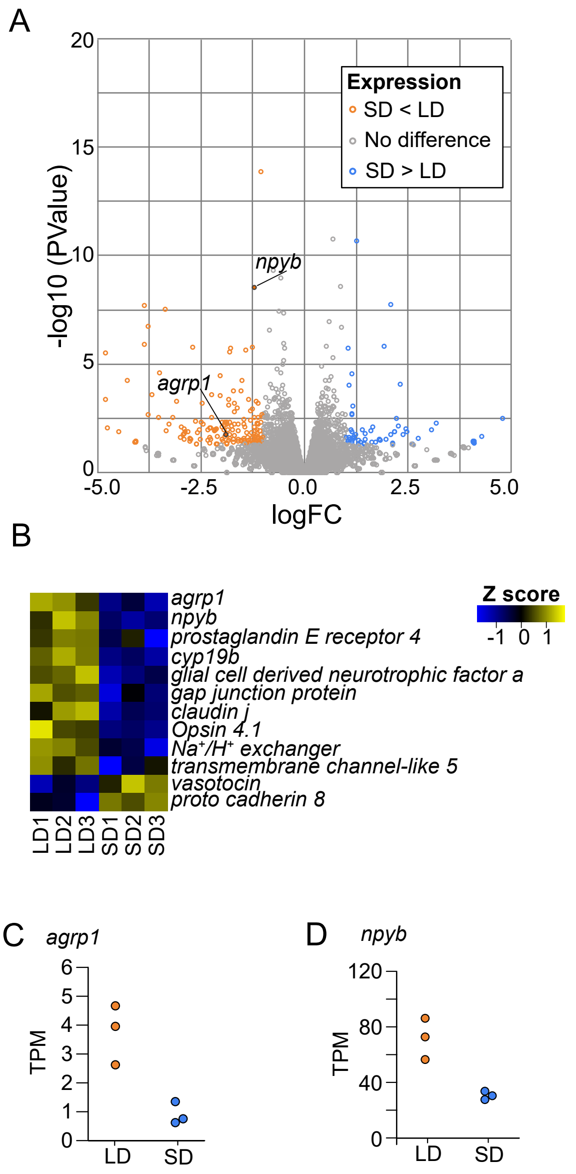
A
-log10 (PValue)
logFC
npyb
agrp1
Expression
SD < LD
No difference
SD > LD
B
Z score
-1 0 1
agrp1
npyb
prostaglandin E receptor 4
cyp19b
glial cell derived neurotrophic factor a
gap junction protein
claudin j
Opsin 4.1
Na+/H+ exchanger
transmembrane channel-like 5
vasotocin
proto cadherin 8
LD1 LD2 LD3 SD1 SD2 SD3
C agrp1
TPM
LD SD
D npyb
TPM
LD SD

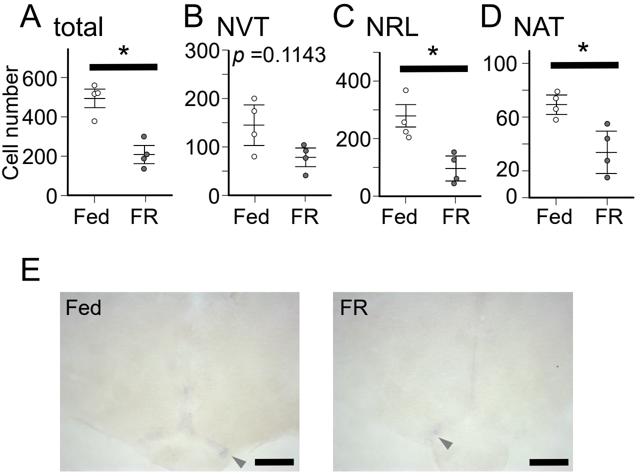
A total
B NVT
C NRL
D NAT
Cell number
*
p =0.1143
Fed
FR
E
Fed
FR

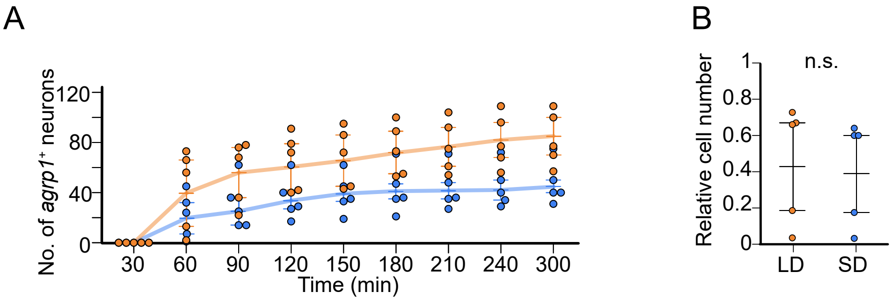

A
No. of agrp1⁺ neurons
120
80
40
0
30
60
90
120
150
180
210
240
300
Time (min)
B
Relative cell number
1
0.8
0.6
0.4
0.2
0
n.s.
LD
SD

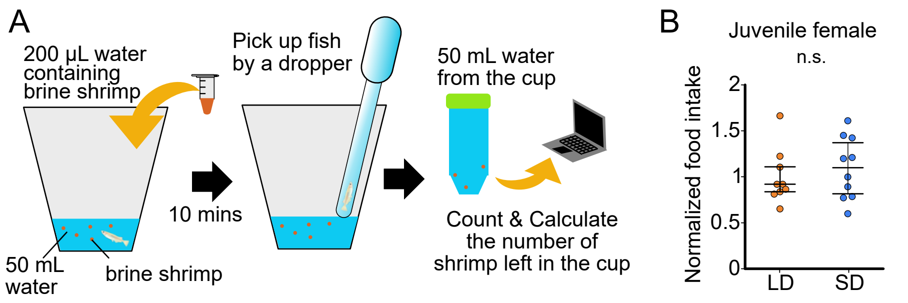
A
200 µL water containing brine shrimp
Pick up fish by a dropper
50 mL water from the cup
10 mins
50 mL water
brine shrimp
Count & Calculate the number of shrimp left in the cup
B
Juvenile female
n.s.
Normalized food intake
2
1.5
1
0.5
0
LD
SD

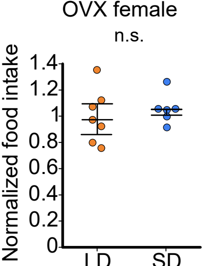
OVX female
n.s.
Normalized food intake
1.4
1.2
1
0.8
0.6
0.4
0.2
0
LD
SD

### A exon 3

***agrp1***$^{+/+}$
GACTCGTCTGCAGGCCTGCAGCTGCAGGGCCGGGCCATGC
GCTCGTCTCGCCGCTGCATCCCTCACCAGCAGTCCTGCCTG
GGTTACCCTCTGCCCTGCTGTGACCCCTGCGACACCTGTTAT
TGCCGCTTCTTCAATGCCATCTGTTACTGCCGCCAAATAGGC
CACACCTGTGCACCCCGGCGCACCTAA

***agrp1***$^{-/-}$
GACTCGTCTGCAGGCCTGCAGCTGCAGGGCCGGGCCATGC
GCTCGTCTCGCCGCTGCATCCCTCACCAGCAGTCCTTGCCT
GGGTTACCCTCTGCCCTGCTGTGACCCCTGCGACACCTGTT
ATTGCCGCTTCTTCAATGCCATCTGTTACTGCCGCCAAATAG
GCCACACCTGTGCACCCCGGCGCACCTAA

### B Protein

***agrp1***$^{+/+}$
DSSAGLQLQGRAMRSSRRCIPHQQSCLGYPLPCCDPCDTCYC
RFFNAICYCRQIGHTCAPRRT

***agrp1***$^{-/-}$
DSSAGLQLQGRAMRSSRRCIPHQQSLPGLPSALL*

## C

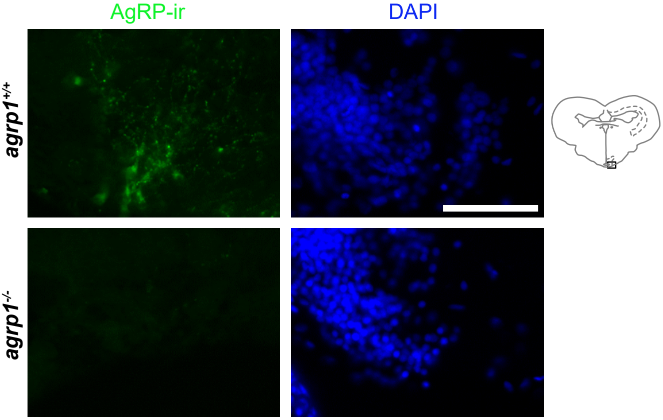

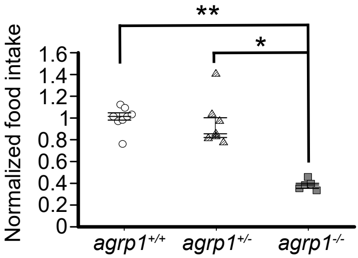
**
*
Normalized food intake
1.6
1.4
1.2
1
0.8
0.6
0.4
0.2
0
agrp1+/+
agrp1+/-
agrp1-/-
